# Single cell multiomics reveal clonal and functional dynamics of MDS stem/progenitor cells during hypomethylating therapy

**DOI:** 10.64898/2026.03.13.711488

**Authors:** Julie A.I. Thoms, Henry R. Hampton, Priscilla L.S. Boon, Olivia Stonehouse, Xiaoheng Zou, Hoi Man Chung, Forrest C. Koch, Feng Yan, Swapna Joshi, Mary N.T. Nguyen, Dorothy Hung, Dale C. Wright, Fatemeh Vafaee, Mark N. Polizzotto, Alexander Swarbrick, Yuanhua Huang, Christopher J. Jolly, Fabio Zanini, John E. Pimanda

## Abstract

Progressive somatic mutations in hematopoietic stem cells (HSCs) drive the development of myelodysplastic neoplasms (MDS). Hypomethylating agents such as azacitidine (AZA) can improve blood counts and reduce blasts, although responses are rarely durable. Determinants of AZA response are complex and incompletely understood, although accumulating evidence suggests that epigenetic rewiring of mutated HSCs underlies improved hematopoietic output. Using single cell multiomics on longitudinal bone marrow samples, we show that AZA responsiveness involves expansion of cells with transcriptomic profiles shared with hematopoietic stem and progenitor cells (HSPCs) from healthy donors. These regenerating cells are depleted of copy number variations and of *TP53* mutations. We also identify patient-restricted cell populations, some of which recede through transcriptional restoration or AZA cytotoxicity, and others which expand, regardless of initial clinical response, and dominate at progression. Individual patients carried multiple patient-restricted populations which had unique surface immunophenotypes and were genetically distinct. Strikingly, sorted cells from in vivo progression clones that were AZA-refractive in patients regained AZA-sensitivity when cultured *in vitro*, suggesting that lack of AZA response at the cellular level can be modulated by cell-extrinsic factors *in vivo*. Overall, we find that AZA response involves partial hematopoietic regeneration via functional differentiation of mutated, but not cytogenetically abnormal HSPCs, and that persistence of AZA-refractive sub-populations contributes to eventual disease progression.

## Introduction

Myelodysplastic neoplasms (MDS) are a heterogenous group of malignancies that are part of a spectrum of disease entities including clonal hematopoiesis of indeterminate potential (CHIP), which mostly occur in older individuals and are characterized by somatic mutations in hematopoietic stem cells (HSCs) and increased risk of progression to acute myeloid leukemia (AML) ^1–8^. Age-associated sterile inflammation contributes to these diseases, with mutations in HSCs thought to confer clonal advantage in an inflamed BM microenvironment ^9–12^. Higher-risk (HR) MDS patients who are ineligible for allogeneic bone marrow (BM) transplant are treated using hypomethylating agents (HMA) including azacitidine (AZA), with around half of patients achieving clinical response characterised by improved peripheral cell counts and delayed progression to AML ^13,14^. The cellular determinants of AZA response are complex and only partially explained by the mutational spectrum ^15–18^. We and others have demonstrated that, with the exception of *TP53* mutated cells ^19^, clinical response is largely driven by improved output from mutated HSCs rather than eradication of mutated cells ^19–24^. However, disease progression typically follows initial response, and whether clinical benefit arises from AZA-induced regeneration of all mutated cells, or only a fraction of subclones have capacity to transiently restore hematopoiesis, remains unknown.

To address these questions and characterize the single cell landscape of AZA-treated MDS, we applied cellular indexing of transcriptomes and epitopes by sequencing (CITEseq) to pre- and post-treatment BM HSPCs from patients uniformly treated with AZA in a clinical trial ^25^. AZA response involved partial hematopoietic regeneration, although cytogenetically abnormal HSPCs did not contribute. However, persistence of AZA-refractive populations contributed to eventual disease progression.

## Results

To understand single cell features of HR-MDS, we performed CITEseq on CD34^+^ cells collected longitudinally from AZA-treated patients on a clinical trial ^25^ (NCT03493646; Figure 1a, Supplemental Figure 1a, b, Supplemental Table 1). To minimise batch effects, cells from each patient/timepoint were spread over multiple captures (Supplemental Figure 1c). Transcriptomic (Tx) profiles from all patients, together with those from five healthy donors, were combined into a UMAP embedding based on Tx features (Supplemental Figure 1d). Clustering of the data assigned 26 Tx clusters (Figure 1b). Cluster 25 was very small (26 cells, remaining clusters range from 186-3667 cells) and not considered for further analyses.

**Figure 1:**
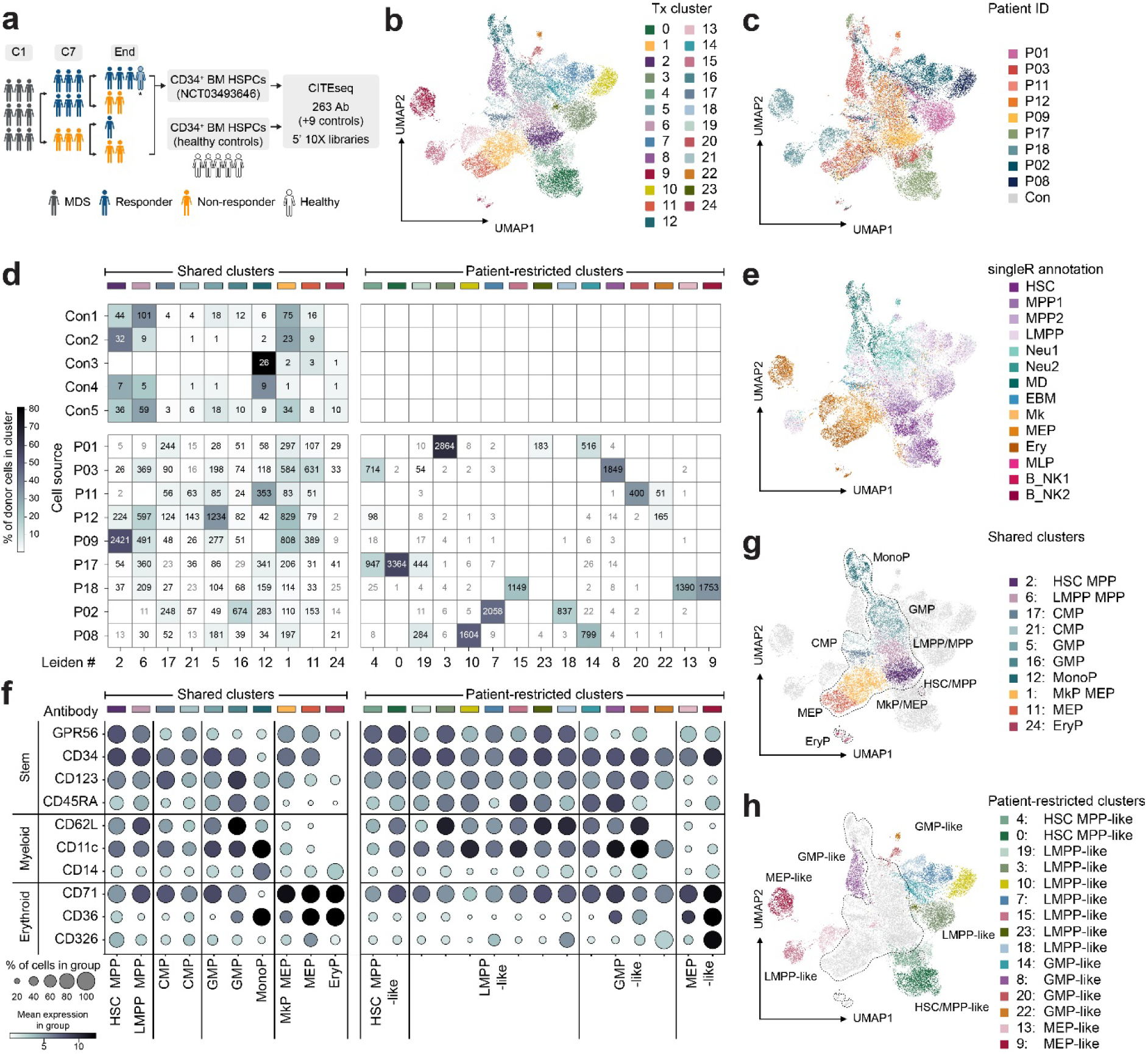
MDS patient-derived HSPCs contain transcriptomic clusters shared with healthy donors along with patient-restricted transcriptomic clusters. **a)** Schematic showing HR-MDS patients and healthy donors. Patient samples were collected at diagnosis (C1), after 6 treatment cycles (C7), and at end of trial (after 12 cycles or at progression). Blue indicates clinical response (complete response [CR], marrow CR [mCR]), orange indicates non-response (stable disease [SD], AML Progression). * insufficient cells for analysis at C12. **b)** UMAP embedding colored by leiden cluster assignment. **c)** UMAP embedding colored by patient-derived and control cells. **d)** Distribution of cells across clusters. Heatmap shows cell distribution between clusters for each patient/donor, with the total cell number in each cluster indicated. White squares with grey text mark clusters containing < 0.5% of patient/donor cells. **e)** UMAP embedding colored by SingleR cell type annotations. **f)** CITEseq antibody tag dotplot for selected antibodies characteristic of stem, myeloid, and erythroid HSPCs. **g)** UMAP embedding colored by shared clusters. **h)** UMAP embedding colored by patient-restricted clusters. Dashed lines in **g)** and **h)** indicate 95% kernel density contours of UMAP coordinates for the shared clusters.

Transcriptomic clusters contained cells from multiple captures (Supplemental Figure 1e). Ten clusters contained cells from patients and healthy donors (Figure 1c, d, Supplemental Figure 1f) and were annotated as “shared” clusters to reflect the commonality of their transcriptional programs. The remaining clusters predominantly contained cells from one, or less frequently two, patients and did not contain cells from healthy donors. We annotated these as “patient-restricted” clusters. To further annotate clusters, we mapped the transcriptome of each cell to a reference which included older individuals ^26^ (Figure 1e, Supplemental Figure 1g) and inspected CITEseq data from key surface markers ^27^ (Figure 1f). Shared clusters mapped to the centre of the UMAP projection with clusters resembling stem cells diverging to myeloid and erythroid branches (Figure 1g). Cells in shared clusters were therefore annotated using conventional HSPC sub-population nomenclature (haematopoietic stem cell/multipotent progenitor (HSC/MPP), lympho-myeloid primed progenitor (LMPP)/MPP, common myeloid progenitor (CMP), granulocyte monocyte progenitor (GMP), monocyte progenitor (MonoP), megakaryocyte erythroid progenitor/megakaryocyte progenitor (MEP/MkP), MEP, erythroid progenitor (EryP). Patient-restricted clusters were generally located toward the periphery of the UMAP (Figure 1h). While these clusters have broad transcriptomic similarity to HSC/MPP, LMPP, GMP, or MEP (Supplemental Figure 1g), to emphasise that these are distinct from healthy-donor-derived cells the assigned cluster (CL) number was used as their primary annotation. Overall, the CD34^+^ compartment in HR-MDS patients contained diverse cell types, some resembling healthy HSPCs and others with unique transcriptomic and immunophenotypic features distinct from healthy donor cells.

We next looked at temporal dynamics of BM CD34^+^ cells over the course of AZA treatment (Supplemental Figure 2). Cluster abundance varied over time in all patients. For example, in P03 (responder at C7 and C12) CL8 became less abundant over time while CL4 expanded; in P08 (non-responder at C7 followed by Progression) CL14 was small at diagnosis, expanded at C7, and then contracted while CL10 expanded. A striking feature of the temporal variations within each patient is the emergence of cells in shared clusters at clinical response timepoints (Figure 2a, b, c) but not at non-response timepoints (Figure 2 b, c, d). Response was associated with an increased proportion of cells belonging to shared clusters and reduction in cells belonging to patient-restricted clusters (Figure 2e). Two patients who responded to AZA and then progressed (P17 and P18) had increased shared cluster cells at the response point which were replaced by patient-restricted cells at AML progression. For two patients (P12 and P09), most cells at diagnosis belonged to shared clusters, albeit with skewed abundance (high GMP in P12 and high HSC/MPP in P09). The over-represented population was reduced at the response timepoints with concomitant increase in erythroid-biased progenitors. We calculated a diversity index for each patient and timepoint which captures the proportion of cells from shared clusters and the relative abundance of patient-restricted clusters; cellular diversity generally increased at response timepoints (Figure 2e).

**Figure 2:**
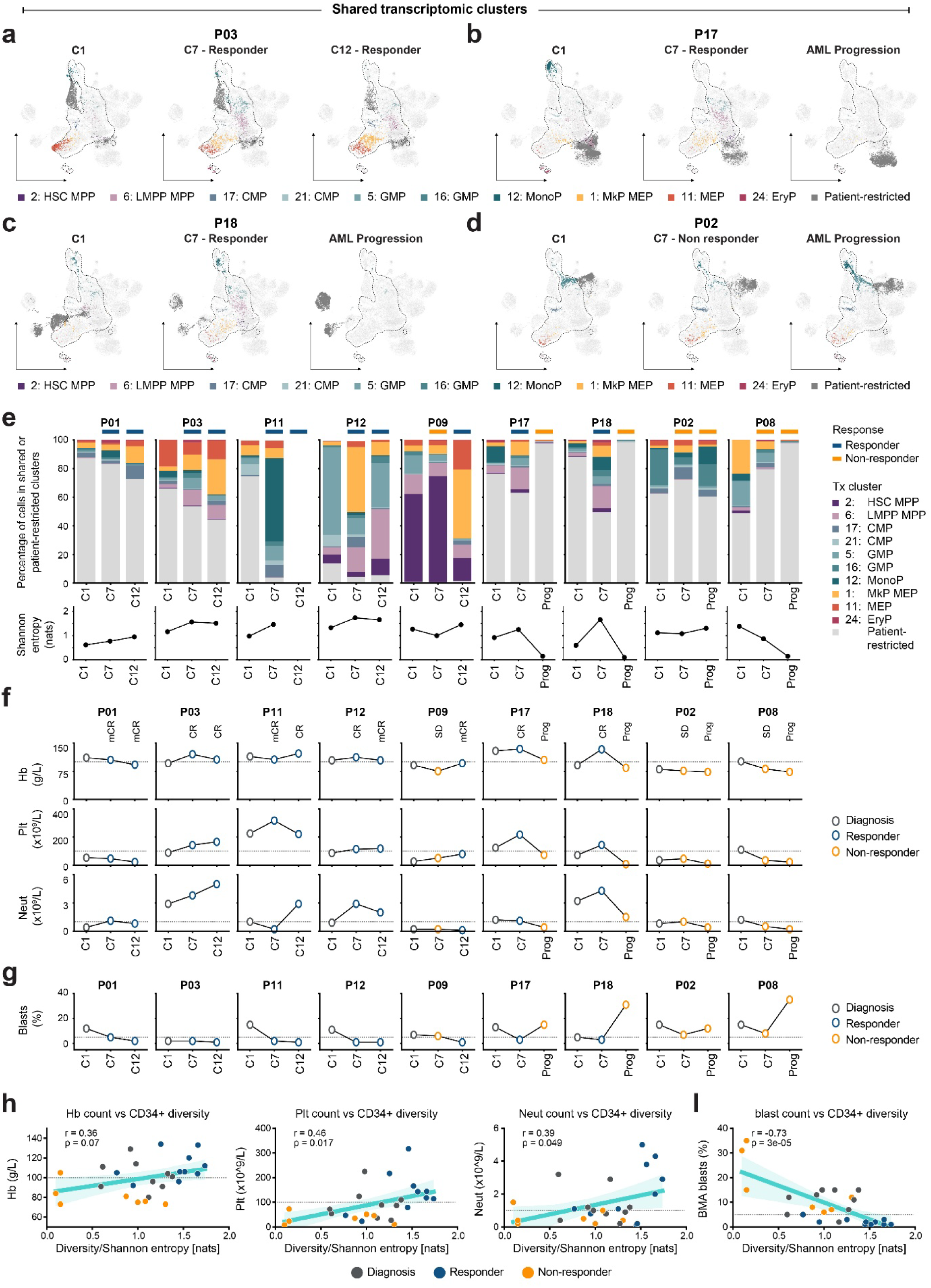
Clinical response to AZA is characterised by increased abundance of HSPC clusters shared with healthy donors. **a-d)** UMAP embeddings showing cell distribution at all time points for 4 example patients with varying clinical trajectories. Dashed lines indicate 95% kernel density contours of UMAP coordinates for the shared clusters. **e)** Bar plots showing relative proportions of shared cell types in all patients at all time points, and line graphs showing cellular diversity (Shannon entropy) at over time. **f)** Peripheral blood (haemoglobin (Hb), platelets (Plt), and neutrophils (Neut)) and **g)** bone marrow (blast) counts corresponding to time points shown in **e)**. Dashed line indicates clinical cut off to call response (Hb: 100g/L, Plt: 100 x 10^9^/L, Neut: 1 x 10^9^/L, Blasts: 5%). **h-i)** Pearson correlations between cellular diversity (Shannon entropy) and blood cell counts shown in **f-g)**.

Single cell changes in CD34^+^ cells reflected clinical observations (Figure 2f, g). Over the whole cohort, increased diversity in the CD34^+^ compartment was positively correlated with peripheral haemoglobin (Hb), platelet (Plt) and neutrophil (Neut) counts (Figure 2h), and negatively correlated with BM blasts (Figure 2i). Overall, AZA-response is accompanied by increased cellular diversity in the BM CD34^+^ compartment, likely as a direct causal relationship since emerging shared clusters contain precursors of circulating cells.

A different pattern was evident in the dynamics of patient-restricted clusters (Figure 3a, b, c, d, Supplemental Figure 2). Total patient-restricted clusters generally reduced in abundance at response points (Figure 3e). Some patient-restricted clusters receded during AZA treatment, even in patients who did not demonstrate clinical response (e.g. CL18 in P02); conversely, expansion of patient-derived clusters during AZA treatment was observed in some patients who showed clinical response (e.g. CL4 in P03). Some patient-restricted clusters underwent substantial expansion, especially at AML progression (e.g. CL0 in P17). Cells belonging to highly expanding clusters were generally detectable even at early time points, including when the patient showed clinical improvement (e.g. CL9 in P18). Some clusters had complex expansion patterns, for example CL14 in P08 which initially expanded, but later receded in favor of CL10 which was present at diagnosis but did not initially expand. Overall, the patient-restricted clusters often persisted throughout AZA treatment; such clusters appeared to evolve during treatment, with minor clusters at diagnosis frequently expanding to become the dominant cluster at progression.

**Figure 3:**
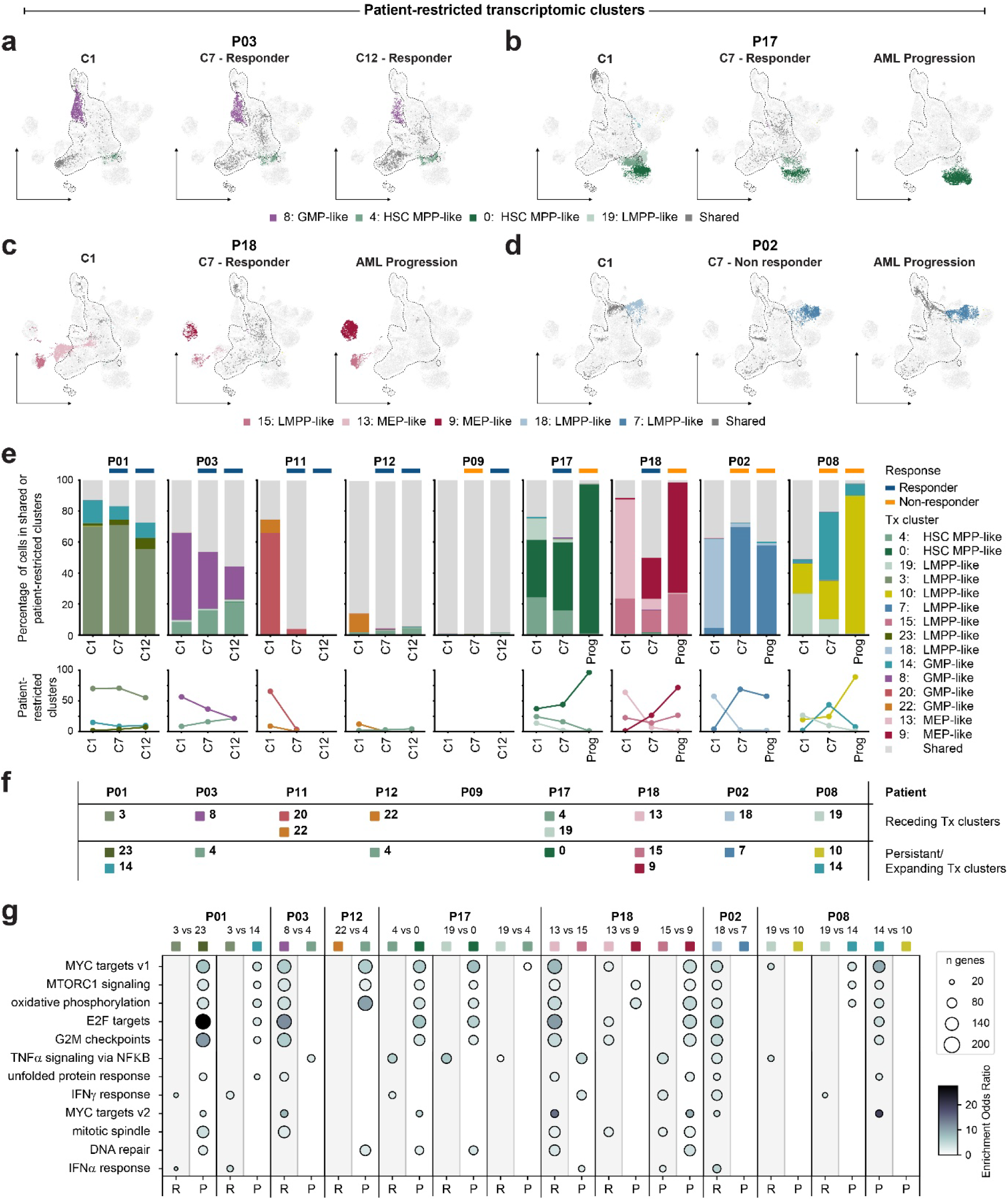
Patient-restricted CD34^+^ clusters persist, recede, and expand during AZA treatment. **a-d)** UMAP embeddings showing cell distribution at all time points for 4 example patients with varying clinical trajectories. Dashed lines indicate 95% kernel density contours of UMAP coordinates for the shared clusters. **e)** Bar plots showing relative proportions of patient-restricted cell types in all patients at all time points (upper) and changes in cluster abundance over time (lower). **f)** Receding and persistent/expanding clusters from each patient. **g)** Hallmark pathway enrichment in genes differentially expressed between receding (R) and persisting/expanding (P) clusters or where present (P18, P08), between two persisting/expanding clusters with different temporal dynamics. The 12 most commonly enriched pathways are shown.

To better understand determinants of cluster dynamics, we performed differential gene expression in clusters that persisted or expanded compared to clusters that receded during AZA treatment (Figure 3f) and looked for enrichment of MSigDB hallmark gene sets ^28^. Twelve pathways were significantly enriched in at least five receding/expanding pairs, with DNA repair, and oxidative phosphorylation commonly enriched in persisting/expanding clusters and immune-related pathways such as IFNγ response, TNFα signalling via NFκB, and IFNα response commonly enriched in receding (Figure 3g).

Patient-restricted clusters might be distinct genetic clones or could represent alternate epigenetic states with altered sensitivity to AZA treatment. Single cell analysis can resolve complex cellular hierarchies in myeloid malignancies ^23,29–31^. We leveraged karyotypic abnormalities in three patients in our cohort (P18, P17, P03) to investigate clonal dynamics within those individuals. Copy number variation (CNV) in individual cells can be inferred based on chromosomally binned gene expression (inferCNV) profiles ^32^. InferCNV at diagnosis revealed cells with widespread karyotypic abnormalities in all 3 patients (Figure 4a, b, c); these patterns broadly matched expected karyotypes (Supplemental Table 1). In P18, the two patient-restricted clusters observed at diagnosis (CL13 and CL15) had similar CNV profiles (Figure 4a). In P17, chr 5q deletion was a common feature in all patient-restricted clusters, but cells in the three persisting/expanding clusters had distinct CNV patterns with CL0 harbouring deletions in chr 7 and chr 12, while CL19 had deletions in chr 17 and CL19 and CL4 both had amplifications in chr 8. Some of the CNV features shared between CL19 and CL4 (e.g. chr 5q del) were also observed in a subset of cells from shared clusters, mostly in cells annotated as LMPP/MPPs (Figure 4b). In P03 many cells (including both patient-restricted clusters and a large MEP cluster) carried a 5q deletion (Figure 4c).

**Figure 4:**
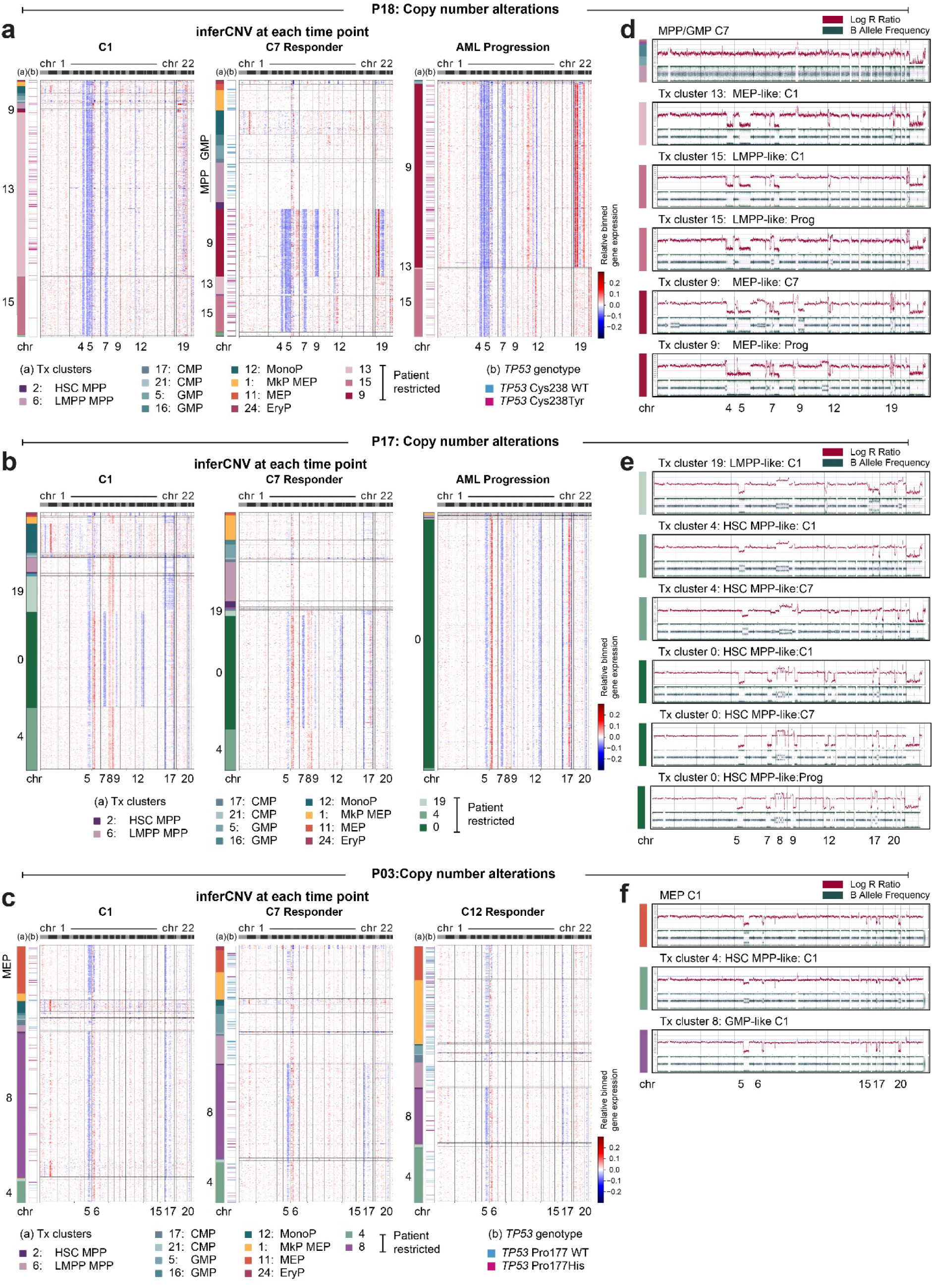
Transcriptomically-defined patient-restricted clusters are cytogenetically distinct. **a-c)** inferCNV heatmaps at each timepoint for **a)** P18, **b)** P17, **c)** P03. Blue colour indicates genomic regions with relatively low gene expression suggesting chromosomal deletions while red colour indicates genomic regions with relatively high gene expression suggesting chromosomal amplifications. Annotation bars indicate (a) transcriptomic (Tx) cluster, and (b) *TP53* genotype (P18 and P03 only). **d-f)** Copy number variation (CNV) in sorted transcriptomic clusters for **d)** P18, **d)** P17, **d)** P03. Red (upper) plot shows Log R Ratio (indicating copy number changes) and green (lower) plot shows B allele frequency (indicating allelic ratio).

All three of these patients experienced an initial response to AZA treatment, accompanied by striking changes in the inferred CNV profiles (Figure 4a, b, c). Cells from shared clusters which emerged at the initial response timepoint (C7) were depleted of CNV abnormalities, suggesting that cells driving clinical response are derived from a residual karyotypically normal pool. However, cells from patient-restricted clusters remained. P18 and P17 subsequently progressed to AML, accompanied by depletion of karyotypically normal cells, and expansion of patient-restricted clusters. Together these data suggest that complex cellular dynamics occur in the CD34^+^ pool during AZA treatment with AZA-induced recovery of productive HSPCs occurring alongside ongoing clonal evolution.

We have previously reported that increased circulating cells following AZA treatment are often derived from highly mutated stem cells, and that eradication of mutated HSC clones is not required for AZA response ^19^. A notable exception was patients carrying *TP53* mutations, where the *TP53* variant (but not other variants in the same individual) was depleted in the circulating cells ^19^. Two of the three patients (P18, P03) with CNV abnormalities in this cohort also carry *TP53* variants, while the third (P17) carries an *ATM* variant; all these variants are predicted to be drivers of the observed abnormal karyotypes. To further investigate the relationship between *TP53* variants and abnormal karyotypes we retrospectively genotyped *TP53* in single cells from our 10X cDNA (the *ATM* variant at amino acid 960 was unable to be recovered). *TP*53 mutation (pink bars) correlated with abnormal CNV profile, with WT *TP53* (blue bars) predominantly found in the karyotypically normal cells emerging during AZA response (Figure 4a, c, Supplemental Table 2).

Inferred CNV data suggested that the patient-restricted clusters are karyotypically distinct; however, inferCNV might misinterpret alternate transcriptomic states as karyotypic differences. For orthogonal validation, we used our CITEseq data to predict surface immunophenotype of Tx clusters, flow-sorted corresponding cell pools (Supplemental Figure 3a, b, c), and karyotyped their nuclear DNA using single nucleotide polymorphism chromosome microarrays (SNP-CMA; Figure 4d, e, f, Supplemental Table 3). For P18, we sorted cells from CL13 and CL15 at diagnosis. SNP-CMA profiles for these populations were consistent with inferCNV data, including chr 19 alterations observed only in CL13 (Figure 4d). CL9 cells sorted at C7 or Progression showed the karyotypic features predicted by inferCNV (including the chr 9 deletion which was observed only at C7), as did CL15 sorted at Progression (Figure 4d). Significantly, sorted cells from shared clusters at the C7 response timepoint did not have detectable karyotypic abnormalities (Figure 4d), confirming our finding that emerging HSPCs which drive clinical response are karyotypically normal.

From P17 we sorted cells from CL0, CL4, and CL19 at diagnosis, and CL0 at progression (Figure 4e). SNP-CMA profiles mostly supported the inferred CNV data, although some inferCNV signal at chr 6 appeared to be due to a transcriptomic change rather than a genuine CNV (Figure 4e). Similarly, we note an inferCNV gain in chr 1 in MonoP populations; although we did not sort this population, the presence of the signal in the MonoP population of all three patients suggests it is due to increased transcription from this region in MonoP cells.

Very few cells were available for P03, but we were able to isolate cells from CL8, CL4, and a population of MEPs at diagnosis (Figure 4f). Although these cells had transcriptomes resembling MEPs, many cells in this cluster carried the *TP53* variant and had karyotypically abnormal inferCNV profiles. As for the previous 2 patients, SNP-CMA profiles confirmed the inferCNV findings (Figure 4f). Overall, sorted Tx clusters have SNP-CMA profiles which essentially match inferCNV profiles, providing orthogonal validation of inferred karyotypes.

To further elucidate clonal relationships between Tx clusters in P18, P17, and P03, we performed UMAP embedding and clustering of cells based entirely on inferCNV profiles (Supplemental Figure 4a, b, c) then combined clustering and timepoint annotations for visualization (Figure 5a, b, c). Nine inferCNV clusters emerged for P18. Cells belonging to shared transcriptomic (Tx) clusters were predominantly in a single inferCNV cluster, except for MonoP cells, while the 3 patient-derived Tx clusters were spread across 6 inferCNV clusters (Figure 5a). Eight inferCNV clusters emerged for P17. Cells from shared Tx clusters were again located in two inferCNV clusters, while the 3 patient-derived Tx clusters were spread across 5 inferCNV clusters (Figure 5b). Only two inferCNV clusters emerged for P03, and these did not perfectly correlate with Tx clusters. Notably, Tx CL4 was split across both inferCNV clusters at all timepoints, consistent with *TP53* genotyping which also revealed a mixture of WT and MT cells in CL4 (Figure 5c). CL4 was one of the few patient-restricted clusters containing cells from multiple patients (P03, P17, P12). This suggests the intriguing possibility that some abnormal cell states, such as CL4, can be accessed via multiple routes including, but not limited to, karyotypic alterations.

**Figure 5:**
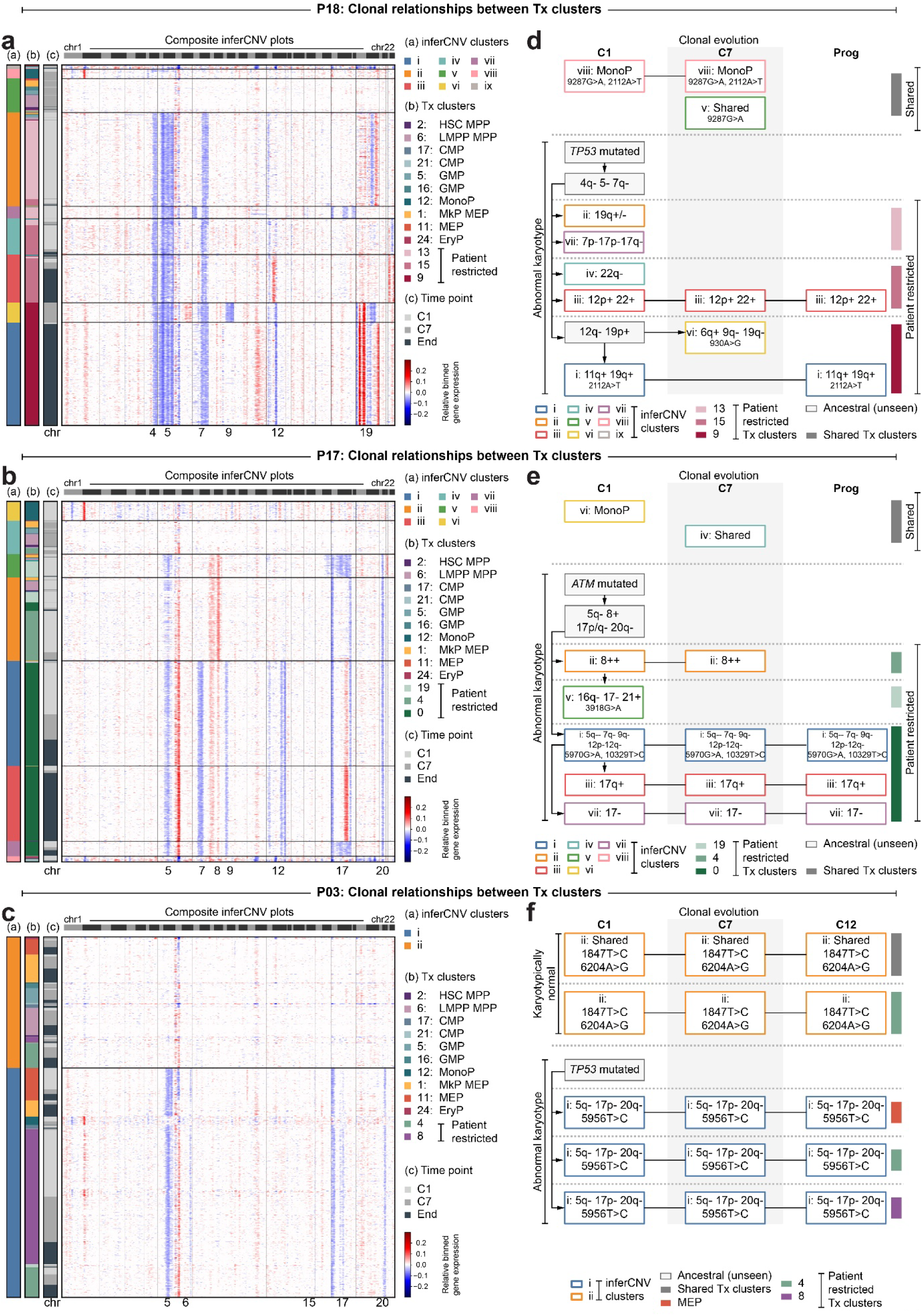
Clonal relationships between transcriptomically-defined patient-restricted clusters. **a-c)** Clustered inferCNV heatmap. Left hand bars show (a) inferCNV clusters, (b) transcriptomic clusters, and (c) timepoint for **a)** P18, **b)** P17, **c)** P03. **d-f)** Inferred subclonal structure with CNV and mitochondrial variations for **d)** P18, **e)** P17, **f)** P03. inferCNV cluster ix not included in phylogeny for P18. inferCNV cluster viii not included in phylogeny for P17.

We also tracked variations in mitochondrial transcripts which could provide increased resolution of historical clonal branchpoints. Mitochondrial variation is sparsely captured from 10X data, nonetheless, we identified informative sequence variants, finding 11 variants in P18, 10 in P17, and 19 in P03 (Supplemental Figure 4di, ei, fi). Clustering revealed combinations of mitochondrial variants corresponding to specific inferCNV clusters (Supplemental Figure 4dii, eii, fii). We then mapped similarity between inferCNV and mitochondrial variant based clusters (Supplemental Figure 4diii, eiii, fiii).

Combining inferCNV based clustering and mitochondrial variations, we constructed clonal phylogenies for P18, P17, and P03 (Figure 5d, e, f). Complex clonal phylogenies with parallel evolutionary events were apparent for P18 and P17. In P18, we inferred the presence of a common ancestral clone with *TP53* mutation followed by loss of chr 4q, chr 5, and chr 7q. Multiple daughter lineages were apparent, and Tx clusters contained multiple genetic subclones, all present at diagnosis except for inferCNV cluster vi (CL9) which was transiently observed at C7 (Figure 5d). In P17 we inferred a common ancestral clone with loss of chr 5q and gain of chr 8, likely derived from an initiating *ATM* mutated cell. Once again, multiple daughter lineages were present at diagnosis with little subsequent clonal evolution (Figure 5e). Clonal structure in P03 was less complex, reflecting the clinical trajectory of this patient who had a sustained clinical response to AZA treatment (Figure 5f). Overall, we find substantial evidence indicating that at least in patients with abnormal karyotypes, the Tx clusters we observe are distinct genetic clones rather than alternate epigenetic states with differing responses to AZA treatment.

To test whether the in vivo differences in AZA response between genetic subclones was due to direct AZA effects on CD34^+^ cells or mediated by other cell types in the bone marrow we used an established *in vitro* stromal co-culture system ^20^ to compare AZA sensitivity between MDS subclones and healthy cord blood (CB) cells. CD34^+^ cells corresponding to specific Tx clusters were sorted at either diagnosis or AML progression (Figure 6ai, bi) or in the case of P17 CL0, at both timepoints.

**Figure 6:**
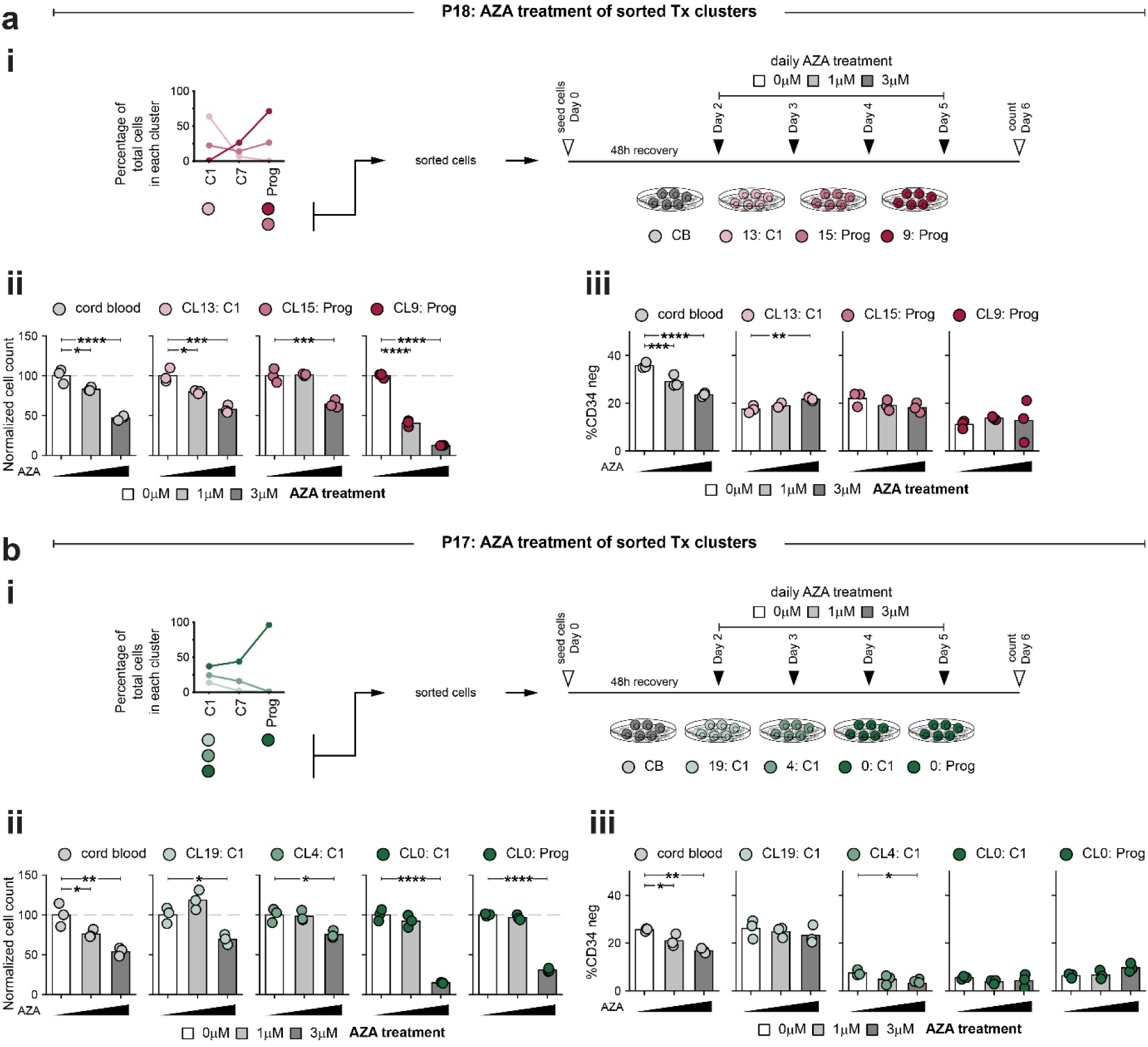
Cells from sorted transcriptomic (Tx) clusters have distinct *in vitro* properties. **a-b)** *in vitro* culture or sorted transcriptomic clusters. **i)** Schematic showing *in vitro* co-culture assay and the sorted populations used for co-culture. **ii)** Proportion of total cells recovered at end of experiment, normalised to 0µM control. Grey dashed line indicates 100%. **iii)** Proportion of total recovered cells which are CD34^-^ (ie/ cells which have differentiated in culture) for **a)** P18, **b)** P17. Statistical comparisons are one-way ANOVA with Tukey’s test for multiple comparisons; only comparisons to vehicle are shown. * indicates *P*<0.05, ** *P*<0.01, \*\*\**P*<0.001, \*\*\*\**P*<0.0001.

CB cells were moderately sensitive to AZA treatment (IC_50_ ∼3µM, Figure 6aii, bii) while sorted MDS cells had variable sensitivity. CL13 and CL15 (P18) had similar AZA sensitivity to CB cells, although CL15 was less sensitive to 1µM AZA (Figure 6aii). CL9 was far more sensitive to AZA *in vitro* than CB cells (estimated IC50 < 1µM), a surprising finding since this cluster was the dominant progression clone. P17 cells from all clusters had less *in vitro* AZA sensitivity than CB cells at the 1µM AZA concentration. For CL19 and CL4 this relative resistance extended to the 3µM concentration (Figure 6bii). However, the dominant cluster at AML progression (CL0) showed increased *in vitro* AZA sensitivity in both diagnosis and progression samples (Figure 6bii).

We used loss of surface CD34 expression at endpoint in co-culture assays as an *in vitro* surrogate for differentiation capacity. CB cells undergo substantial differentiation in culture, which was slightly attenuated in the presence of AZA (Figure 6aiii, biii). Cells from P18 had somewhat lower differentiation capacity, and the progression cluster (CL0) showed the lowest proportion of CD34^-^ cells following AZA treatment. Cells from P17 CL19 had similar differentiation capacity to CB cells, but the CL4 and CL0 cultures had very low endpoint proportions of CD34^-^ cells at all AZA concentrations. Overall, sorted MDS clusters have diverse *in vitro* responses to AZA, consistent with their identity as distinct genetic clones, and where tested, cluster properties were maintained over a period of months within the patient (P17 CL0). However, clusters which dominated at progression in patients were the most AZA-sensitive *in vitro*, suggesting that clones which expand in the patient context may still maintain intrinsic AZA sensitivity.

## Discussion

Here we applied single cell multiomics to longitudinal bone marrow CD34^+^ samples and showed that AZA-response involves expansion of cells with transcriptomic profiles shared with HSPCs from healthy donors. Regenerating cells were depleted of copy number variations and of *TP53* mutations. We also identified patient-restricted cell populations, some of which expanded, including during periods of clinical response, and dominated at AML progression. Patient-restricted cell populations could be isolated based on unique immunophenotypes and have potential utility in tracking measurable residual disease (MRD) and *ex vivo* drug testing.

The origin of the stem cell pool which underpins HMA response is an open question. We and others have demonstrated that AZA response does not require ablation of mutated cells ^19–22^, which, with the exception of *TP53* mutated cells, contribute proportionally to increased peripheral counts ^19^. Here we identify CD34^+^ cells with transcriptomic features of healthy cells which emerge during clinical response and disappear at relapse/progression. In karyotypically abnormal MDS, emerging cells originate from an unmutated, karyotypically normal pool. MDS patients with *TP53* mutations have higher frequency of HMA response but these are rarely durable ^33–35^. One possibility is that response in these patients is driven by a residual, perhaps initially quiescent, pool of *TP53* WT cells that are reactivated by AZA treatment. Subsequent relapse may be driven by exhaustion of this pool or alternatively, may reflect rapid outgrowth of the *TP53* mutated clone. In patients lacking karyotypic abnormalities it is unclear whether a residual unmutated pool of stem cells is reactivated and contributes to circulating mature cells, although persistent high VAF frequencies in some patient at CR ^19^ suggest that such cells are rare. If residual unmutated cells are limited or non-existent, and clinical response to HMAs is primarily driven by epigenetic rewiring of mutated cells, then aggressive treatment strategies aimed at eradication of all mutated cells may be counterproductive in the absence of an allograft with healthy HSCs. Also unclear is the extent to which remodelling of the BM microenvironment contributes to HMA response. Inflammatory changes in the bone marrow microenvironment have been extensively implicated in driving clonal selection in myeloid disease ^10–12^. AZA response likely depends not only on cell-intrinsic response in CD34^+^ cells, but also on AZA-dependent changes in the BM microenvironment ^36–38^ which may in turn govern which specific cells are supported to increase productive output, be they residual healthy cells or mutated MDS clones.

Relapse during AZA treatment can be accompanied by changes in sub-clonal structure ^39,40^ which could potentially be used to predict disease progression. Standardised MRD guidelines are not yet established for MDS ^41–43^ and are hindered by potential confounding effects including disease heterogeneity, and the presence of molecular lesions originating from antecedent CHIP which may not indicate impending relapse and are recommended for exclusion from NGS-based MRD measures in AML ^44^. We observed marked distribution changes in Tx-defined CD34^+^ clusters during AZA treatment that may be useful for MRD tracking but present a challenge due to the feasibility of deploying scRNAseq technologies in a diagnostic setting. However, we were able to isolate cells from these Tx clusters based on unique surface immunophenotypes, providing proof of principle that flow cytometry based MRD tracking of patient-specific clusters is possible.

A counterintuitive finding in this study was that MDS subpopulations that expanded during treatment to dominate the CD34^+^ compartment at relapse were profoundly sensitive to AZA in an *ex vivo* co-culture system. While the mechanisms underpinning acquired AZA resistance are incompletely characterized, metabolic rewiring of pyrimidine synthesis pathways, particularly in a TP53 mutant context, is a potential mediator ^45,46^. Culture systems based on conventional cell media designed to support cell proliferation may well be able to override any *in vivo* metabolic rewiring. Alternatively, *in vivo* AZA-sensitivity might be modulated by cell-extrinsic factors such as cell-cell interactions or soluble factors originating from an altered bone marrow niche. Given the high level of heterogeneity in MDS patients, improved treatment outcomes will likely require individualized treatments. Capacity to accurately predict drug responses in *ex vivo* culture systems will facilitate development of such treatments, but culture systems that recapitulate physical and immunological features of the BM milieu will be required. Nonetheless, our *ex vivo* culture data demonstrates that cell populations which expand in patients in the context of AZA treatment retain capacity for AZA-response in some contexts. The challenge will be to recapitulate that context in patients.

The number of samples analyzed here was modest (26 BM samples from 9 patients). Nevertheless, the goal was to understand dynamic changes occurring in the CD34^+^ compartment in a cohort of HR-MDS patients uniformly treated with HMA therapy where responders were CR/mCR. Our data revealed common features of AZA response in these molecularly heterogenous patients but also characteristics of atypical cell types that vary from one individual to the next. The study was also limited by the technical challenge of mapping the complex clonal structures implied by longitudinal VAF data onto transcriptomic data. Overcoming this challenge would further illuminate precisely which subclones of cells can reacquire a shared transcriptomic profile, including in karyotypically normal disease.

Overall, we find that clinical response to AZA in HR-MDS is accompanied by emergence of CD34^+^ cells transcriptionally indistinguishable from healthy cells. While mutated stem cells can contribute to improved peripheral counts ^19,21^, we found that genome instability is a critical barrier to increased hematopoietic capacity, potentially explaining why TP53 mutated clones contribute minimally to hematopoietic regeneration in response to AZA treatment ^19^. Furthermore, cells with patient-restricted transcriptomic profiles persist in the CD34^+^ compartment even during clinical response ^47^. Thus, sustained clinical response likely requires either rewiring of mutated HSPCs or activation of residual unmutated stem cells to restore hematopoietic capacity, and concomitant eradication of AZA-refractive cell clones which resist differentiation or apoptosis and serve as a reservoir for disease progression. While at least some patient-restricted cell populations can be prospectively isolated for ex vivo drug testing, development of advanced culture systems that are able to recapitulate in vivo cell behaviour will be required to enable translation of ex vivo drug sensitivity to individualized treatment options.

## Methods

### Collection and processing of bone marrow samples

Human bone marrow samples (Supplemental Table 1) were collected as part of an open label phase 2 multicentre investigator-initiated trial ^25^ (NCT03493646: Evaluating in Vivo AZA Incorporation in Mononuclear Cells Following Vidaza or CC-486). All participants provided written informed consent for clinical research including genetic testing and collection/storage of human tissue. Cord blood units were provided by Sydney Cord Blood Bank. Use of samples received ethical approval (2022_ETH00727) from the South Eastern Sydney Local Health District Human Research Ethics Committee.

Bone marrow aspirates were diluted five-fold in RPMI 1640 (Thermo Fisher, #21870076), cord blood were diluted two-fold in PBS containing 2mM EDTA, then mononuclear cells (MNCs) were isolated using Lymphoprep (ELITech Group, #1114547). Cells with high CD34 expression cells were purified using anti-CD34 conjugated magnetic beads (Miltenyi Biotec, #130-046-702) using an AutoMACS with program posseld2 (Miltenyi Biotec). Cells with lower CD34 expression were subsequently recovered by passing unbound cells back through the AutoMACS using program possel. Selected cells were frozen in cryovials with fetal bovine serum (FBS; Gibco, #10099-141) + 10% DMSO or CryoBrew (Miltenyi Biotec, #130-109-558) + 10% FBS.

### SNP genotyping

MNC-derived genomic DNA from each patient/donor was extracted using a DNeasy Blood & Tissue Kit (Qiagen #69504) and genotyped for 654,027 markers using the Illumina Infinium Global Screening Array-24 v3.0 (GSA v3.0). This array features clinically relevant research variants and quality control (QC) markers. Each assay was conducted using 200 ng of DNA which was amplified and then enzymatically fragmented according to the manufacturer’s protocols. Fragmented DNA was loaded onto BeadChips (GSA-24v3.0_A1) and hybridized overnight at 48°C. The next day, probe extension and staining were performed using the Tecan/IAPS automated liquid handling system. BeadChips were scanned using the iScan system (Illumina) to generate intensity data (.idat files).

SNP calling and QC reporting were carried out with GenomeStudio® v2.0.5 using the GenCall algorithm and the GRCh38 reference build. Illumina’s QC system includes both sample-dependent and sample-independent controls to evaluate the integrity of each step in the workflow and overall sample quality. Intensity data were converted to genotype call files (.gtc) using the Illumina IAAP Genotyping Command Line Interface (v1.1). The bcftools +gtc2vcf program was used to convert the .gtc files to variant call format (.vcf) files ^48^. The Demuxafy pipeline (3.0.0) was then used was used to preprocess the VCF files for input into Demuxlet ^49 50^.

### Pooling strategy for single cell sequencing

Magnetically isolated CD34+ cells from nine patients (three time points each) and five healthy donors were analysed. Samples were pooled to distribute different timepoints and patients across multiple captures (Supplemental Table 4). CD34 high cells were captured in six batches. Four of these captures (which account for 89% of all cells analysed) included CITEseq staining and contained cells from six patients and two controls. A smaller number of CD34 high cells from two patients were captured in HSPC Pools 5-6 which were stained with hash tagging antibodies. CD34 low cells were pooled along with other bone marrow mononuclear cells from the same patients and captured in six batches (four patients per batch) without CITEseq antibodies (Pools 1, 2, 3, 5, 6, 7).

### Preparation and sequencing of CITEseq libraries

A lyophilized custom panel of 273 TotalSeqC antibodies relevant to immuno-oncology (Biolegend; Supplemental Table 5) was removed from the fridge and left at room temperature for 5 minutes. The antibody panel was spun down at 10,000g for 30 seconds then reconstituted with 52.5µL of cell staining buffer (Biolegend, #420201) and left at room temperature for 5 minutes with vortexing after 2 minutes. This CITEseq master mix was transferred to a 2mL Eppendorf tube and spun down at 14,000g for 10 minutes at 4°C and used immediately to stain prepared cells.

Cells were thawed in a waterbath then transferred to a new tube and washed twice with FACS buffer (PBS + 2% FBS + 1mM EDTA). Live cells were selected by staining with 0.3ng/mL DAPI (BD Biosciences #564907) and sorting the DAPI-negative population into PBS + 5% FBS on an Aria III Fluorescent-Activated-Cell-Sorter (BD Biosciences). Cells were then counted using a haemocytometer. For HSPC Pools 5-6, cells from individual donors were blocked with 2μL of Human TruStain FcX (Biolegend, #422302) in 20μL of FACS buffer for 15 minutes on ice. 1µL of hashtag antibody was added, and the cells incubated on ice for 20 minutes then washed 4 times with 1mL of PBS + 5% FBS at 4°C prior to pooling. For all capture pools, samples were pooled into a single tube with at least 100000 total cells per pool. For HSPC Pools 1-4, cells were blocked with 2μL of Human TruStain FcX (Biolegend, #422302) in 20μL of FACS buffer for 15 minutes on ice. 12.5μL of prepared CITEseq master mix was added to the sample, and the cells incubated on ice for 20 minutes. Cells were washed 3 times with 1mL of PBS + 5% FBS at 4°C. A 40μm FlowMi strainer (Merck, #BAH136800040-50EA) was used to remove cell aggregates. The total volume was then topped up to 3mL with PBS + 5% FBS and cells were spun down at 500g for 7 minutes at 4°C. 25,000 cells were loaded onto a Chromium Next GEM Single Cell 5’ GEM v2.0.

Gene expression and CITEseq libraries were generated according to the manufacturer’s protocols. The resulting libraries were sequenced on an Illumina Novaseq 6000 using the S4 Reagent kit (100 cycles) at the Ramaciotti Centre for Genomics (UNSW Sydney). The libraries were sequenced with a target depth to recover 70,000 reads per cell for the gene expression library and 55,000 reads per cell for the CITEseq library.

### Preparation and sequencing of amplicon libraries for single cell mutation calling

Excess 10X cDNA from HSPC Pools 1-4 that was not required for sequencing was used to determine single cell genotype at known *TP53* mutations using a previously published protocol ^51^. Three TP53 mutations were tracked; chr17:7578400_G>T corresponding to Pro177His in P03, chr17:7577568_C>T corresponding to Cys238Tyr in P18, and chr17:7579369_G>C corresponding to Ser106Arg in P18. PCR primers used for genotyping are listed in Supplemental Table 6. Template-switching oligo (TSO) artifacts from the 5’ 10X Genomics libraries were removed using five cycles of PCR amplification with the following reagents: 0.25µL of cDNA, 2.5µL of 10µM AA0272 primer and 2.5µL of 10µM Bio-AA0273 primer, 25µl of 2x Kapa HiFi Uracil Kit (Roche, #KK2801), 0.5µL of 100x SYBR Green, 0.1µL of 500x ROX and 18.4µL of nuclease free water. After an initial 3-minute melt step at 98°C, PCR cycles were 98°C for 20 seconds, 65°C for 30 seconds, 72°C for 8 minutes. The resulting PCR products were purified using a 0.6 ratio of Ampure beads (Beckman Coulter Australia, #A63880) and eluted in 10µL of TE pH 8.0. Biotinylated molecules were purified using 20µL of Dynabeads Streptavidin Magnetic Beads (ThermoFisher, #60101) according to the manufacturer’s protocol.

Amplification of target genes was performed using rhPCR. The rhPCR master mix contained: 1.25µL of 20µM rhCGA_venus primer, 2.5µL of 20µM biotinylated gene specific primer, 2.5µL of 20x rhPCR buffer, 0.8µL of 25mM dNTPS, 1.25µL of 20mU RNase H2 (diluted in RNase H2 buffer) (IDT, #11-02-12-01), 2µL of 5U/µL Hotstart OneTaq (New England Biolabs, #M0481L), 0.5µL of 100x SYBR Green, 0.1µL of 500x ROX. The PCR protocol was: 95°C for 5 minutes followed by 96°C for 20 seconds, 60°C for 6 minutes, 72°C for 4 minutes for a total of 17 cycles. PCR products were purified using a 0.6 ratio of Ampure beads, eluted in 10µL of TE pH 8.0 and purified using 20µL of Dynabeads Streptavidin Magnetic Beads (ThermoFisher, #60101) according to the manufacturer’s protocol.

Nested PCR was then used to amplify genes of interest. PCR master mix contained: 1.25µL 20µM CGAvenus.PS primer, 1.25µL 20µM of gene specific nested primer, 0.5µL of 100x SYBR Green, 0.1µL of 500x ROX, 25µL of Q5 High-Fidelity 2x Master Mix (New England Biolabs, #M0492S), 22.15µL of water. The PCR protocol was: 98°C for 5 minutes followed by 98°C for 20 seconds, 62°C for 30 seconds, 72°C for 4 minutes for a total of 17 cycles. PCR products were purified using a 0.6 ratio of Ampure beads.

Amplicon libraries derived from each unique 10X capture were pooled into a single tube, and 500 ng of each of these pools was used as input for end prep and barcoding using the Native Barcoding Kit v14 (Oxford Nanopore Technologies, SQK-NBD114.24). Sixty femtomoles of each barcoded sample were pooled for adapter ligation using the SQK-LSK114 kit (Oxford Nanopore Technologies). A total of 75ng of the final adapter-ligated library was loaded onto a PromethION flow cell (PRO114M R10.4.1), which was run for 72 hours. Raw pod5 data were live basecalled in high-accuracy (HAC) mode on the PromethION, via MinKNOW 24.02.19 and Dorado 7.3.11, using the configuration file dna_r10.4.1_e8.2_400bps_5khz_hac_prom.cfg.

### Fluorescence-activated cell sorting

Cryopreserved patient samples (some MNC, some pre-selected CD34+ cells) were rapidly thawed in a 37 °C water bath and diluted in wash solution (2.5% Dextran 40, 2.5% (w/v) human albumin in 0.9% NaCl). MNC samples were further diluted in saline and enriched by density gradient centrifugation using Lymphoprep (StemCell Technologies, 18061) according to the manufacturer’s instructions. MNCs were washed in FACS buffer (2% heat-inactivated FBS, 1% penicillin–streptomycin, 0.5 mM EDTA), Fc-blocked (BD, #564220), and stained with anti-human CD34-APC (BD, #340441). Cell counts and viability were assessed using methylene blue (StemCell Technologies, #07060), and CD34⁺ cells were enriched by magnetic separation using anti-APC microbeads (Miltenyi Biotec, #130-090-855) using a stand magnet and following the manufacturer’s protocol.

CD34⁺ cells were stained with the indicated antibody panels (Supplemental Table 7) for 45 min on ice in the dark. Viability was assessed by adding 7-AAD (Thermo Fisher Scientific, A1310) immediately prior to sorting. Cells were sorted on a BD FACSAria™ III using a 100um nozzle. Single stain bead compensation controls and purity checks were performed as applicable.

### Single nucleotide polymorphism chromosome microarray (SNP-CMA)

DNA was extracted from sorted cells using an AllPrep® DNA/RNA Micro kit (Qiagen #80284) and copy number variations determined using an Illumina Global Diversity Array Cyto v3 (8x 1,800K SNP microarray), according to the manufacturer’s protocols. The microarray mean effective resolution for copy number variation and copy neutral-LOH 5Mb was 0.015Mb and 5Mb, respectively. Microarray data was analysed using Bionano Genomics VIA (v7) software and the GRCh38 reference genome build. Except when otherwise indicated, standard reporting criteria were used: copy number segments >5Mb, focal copy number alteration <5Mb covering a region of interest, copy-neutral loss of heterozygosity (cnLOH) >10Mb, and mosaicism or acquired clonal abnormalities >15% (Supplemental Table 3). Gene count and annotation was based on the NCBI reference sequence (RefSeq) curated genes.

### Ex vivo drug sensitivity

Drug testing was performed using CD34⁺ cord blood cells (controls) or sorted patient-derived cells cocultured with murine MS-5 stromal cells ^20,52^. Each assay was performed in technical triplicates. MS-5 cells were pre-seeded in 24-well plates at least 24 h prior to the addition of primary cells and allowed to reach confluence. Cells were plated in Gärtner’s medium consisting of IMDM (Thermo, #12440053) supplemented with 12.5% heat-inactivated FBS (Gibco, #10099-141), 12.5% heat-inactivated horse serum (Thermo, #16050130), 2 mM L-glutamine (Gibco, #25030-081), 8 µg/mL monothioglycerol (Sigma-Aldrich, #M6145-25ML), 1 µM hydrocortisone (Sigma-Aldrich, #H0888-1G), and 1× Primocin (InvivoGen, #ant-pm-1). Primary cells were allowed to recover for 48 h before being treated daily with 1 µM AZA (Selleck Chemicals, #S1782), 3 µM AZA, or DMSO vehicle control. After four days of treatment, 10,000 counting beads (Thermo, #C36995) were added to each well prior to harvesting the entire well contents by trypsinization (Thermo, #15400054). Cells were Fc-blocked (BD, #564220) and stained with anti-mouse CD105 PE-Cy5 (Biolegend, #120428, 1:160 dilution), anti-human CD34 PE (BD, #555822, 1:20 dilution), anti-human CD45 FITC (BD, #555482, 1:20 dilution), and DAPI (BD, #564907, used at 0.25µg/ml). Flow cytometry was performed on a BD FACSAria™ III, and data were analysed using FlowJo (v10.9.0). Absolute total cell count in each well was calculated as follows:

Absolute total cell count = (Cell count [DAPI-mCD105-CD45+]/ Bead count) × 10,000

### Data visualization and statistical comparisons

Log R Ratio and B Allele Frequency plots were generated using Bionano Genomics VIA (v7 or later), flow sort plots were generated using FlowJo (v10.9.0), and the oncoprint plot was created using pyoncoprint (v0.2.5) ^53^. All other data was formatted and plotted in python (v3.12.3) using pandas (v2.2.2), matplotlib (v3.10.0) and seaborn (v0.13.2). Shannon entropy was calculated with scipy (v1.13.1) using the tool stats.entropy with cells assigned to patient restricted clusters treated as a single category. Pearson correlation was calculated with scipy (v1.13.1) using the tool stats.pearsonr. One-way ANOVA with Tukey’s test for multiple comparisons was performed using GraphPad Prism (v10.6.1).

### Bioinformatic Analyses

Analysis tools were run with default parameters except when indicted otherwise. Analyses were performed using the computational cluster Katana, which is supported by Research Technology Services at UNSW Sydney.

### scRNAseq data processing

The 10X Genomics single-cell libraries were aligned to the GRCh38 human reference genome (refdata-gex-GRCh38-2020-A) using the CellRanger multi workflow (version 7.1.0)^54^ with a dual-modality configuration for gene expression (GEX) and antibody-derived tag (ADT) reads.

Filtered GEX matrices from each library were loaded into python and analysed using scanpy (v1.10.4) ^55^. Demuxlet, run as part of the Demuxafy workflow, was used to demultiplex the sample identity and detect doublets in each droplet ^50^. Cells with a single genotype, as identified by Demuxlet, were retained. Cells expressing fewer than 200 genes and genes detected in fewer than 3 cells were removed. Additional filtering was conducted using pool-specific thresholds (Supplemental Table 8) for number of genes by counts and mitochondrial gene expression. Appropriate thresholds for each pool were determined based on the count distribution and were set to exclude outlier cells. After filtering, all libraries were concatenated and counts per cell were normalized to a total of (1 x 10^4^) followed by log plus one (log1p) transformation. Principal component analysis was performed to reduce the dimensionality of the data. The first 50 principal components were used to calculate the neighbourhood graph of cells. The graph was then embedded in two dimensions and visualized with UMAP.

Clustering of all cells based on transcriptomic features was performed using scanpy.tl.leiden with default parameters (resolution = 1). HSPCs were isolated by selecting leiden clusters which express CD34 and lack CD3, leaving 38231 stem and progenitor cells for analysis. The number of cells analyzed per patient at each timepoint is shown in Supplemental Table 9. Further leiden clustering of HSPCs produced 26 clusters containing between 26 and 3367 cells. Cluster 25, containing 26 cells, was substantially smaller than all other clusters (the next smallest clusters, cluster 24 and cluster 23, contained 186 and 187 cells respectively) and was excluded from further analysis.

ADT data from each library pool was further processed with Muon (v0.1.7) ^56^. ADT counts were first normalized using the denoised and scaled by background (DSB) method with isotype controls ^57^, then individual library pools were concatenated and merged with the filtered and normalised gene expression libraries. Only cell barcodes also detected in the GEX library were retained. Differential antibody expression analysis was conducted using the rank_genes_group function in scanpy (v1.10.4).

### Annotation of leiden clusters based on transcriptomic features

Transcriptomically defined clusters were annotated as “shared” or “patient-restricted” based on presence or absence of cells derived from healthy donors and the proportion of patient samples contributing to those clusters. Shared clusters contained cells originating from both healthy donors and patients: furthermore, they contained cells originating from multiple different patients. Patient-restricted clusters never contained cells from healthy donors and were predominantly composed of cells originating from one or two patients. To predict identity of each individual cell, reference data was download from http://scrna.sklehabc.com/HSPC/, including gene expression count matrix and cell metadata ^26^. Cells were annotated with SingleR (v 2.6.0) using logcounts for both input and reference dataset and used the de.method “wilcox” and labels from “RNA_clusters” ^58^. Cell surface markers highly consistent in defining of stem, myeloid, and erythroid HSPCs that were also present in our CITEseq antibody set were selected from Zhang et al ^27^. Shared clusters were manually annotated using conventional HSPC sub-population nomenclature (haematopoietic stem cell/multipotent progenitor (HSC/MPP), lympho-myeloid primed progenitor (LMPP)/MPP, common myeloid progenitor (CMP), granulocyte monocyte progenitor (GMP), monocyte progenitor (MonoP), megakaryocyte erythroid progenitor/megakaryocyte progenitor (MEP/MkP), MEP, or erythroid progenitor (EryP) based on frequency of predicted cell types and cluster-wide surface immunophenotype. Patient-restricted clusters were manually annotated to HSC/MPP-like, LMPP-like, GMP-like, or MEP-like based on frequency of predicted cell types and cluster-wide surface immunophenotype and in recognition that clusters found in only a few HR-MDS patients are likely to represent abnormal cell types, albeit with similarities to healthy cell types.

### Differential gene expression, pathway enrichment, and copy number variation

Differential gene expression analysis was conducted using the rank_genes_group function in scanpy (v1.10.4). Pathway analysis was conducted using the scanpy.queries.enrich (a wrapper for gprofiler) and Gene Set Enrichment Analysis (GSEA) ^59–61^. Copy number variation analysis was conducted using infercnvpy (v0.6.0) (https://github.com/icbi-lab/infercnvpy) ^32^ to generate chromosomally binned gene expression (inferCNV) profiles, using data from healthy HSPCs as the control cells negative for CNVs.

### Single cell mutation calling from single cells

Oxford Nanopore (ONT) data were processed using a modified version of the *nanoranger* pipeline ^51^ available at https://github.com/ForrestCKoch/nanoranger. The original pipeline was modified so that barcodes must be an exact match to the cell barcodes output by Cell Ranger to be considered for downstream processing.

Mutation calling was performed using scripts developed in-house and available at https://github.com/ForrestCKoch/OralAZA_ONT_mutations. This script assigns a mutation label to each read that overlaps a predefined mutation site: wild-type (WT), mutant-type (MT), or unknown (UN). A UN label indicates the read sequence at the site did not match either the WT or MT allele—important given the higher error rates of ONT reads. Reads with base quality scores below 25 at the mutation site were excluded from analysis. Per-cell mutation counts were aggregated, and cells were classified using predefined criteria. Cells with > 10 wild-type reads and ≥ 98% of total reads mapping to the WT allele were assigned as wild-type. Cells were called mutant if they exceeded a mutation-specific threshold for MT reads and ≥ 30% of their total reads aligned to the mutant allele. The mutation-specific threshold was set based on read data from patients known to be WT for that variant. For P03 (*TP53* Pro177His), the threshold used was 25 MT reads, and for P18, the threshold was 55 MT reads (*TP53* Cys238Tyr). Although chr17:7579369_G>C corresponding to Ser106Arg was detected in bulk VAF analysis for P18 ^25^, the mutated allele was never detected in either nanopore or short read data from P18, even though abundant wild type reads were present. We interpret this to indicate that the Ser106Arg allele, while present, is not expressed.

BAM files produced by Cell Ranger were analysed for the presence of either *TP53* mutation using pysam (v0.22.1). No discrepancies were found for TP53, that is, there were no cells classified as wild type by *nanoranger* that harboured mutant TP53 reads in the Cell Ranger BAM files.

### Clonal tracking and mitochondrial variation

InferCNV clusters were assigned by leiden clustering within scanpy (v1.9.8). To trace mitochondrial variants within individual patients, mitochondrial allele counts were first extracted from scRNA-seq using cellSNP-lite^62^. Mitochondrial SNV clusters were then assigned by MQuad^63^ followed by SNPmanifold^64^ and mitochondrial similarity was computed based on log-odds of the number of cells in each mitochondrial SNV clusters. Phylogenies were manually constructed for P18, P17, and P03 based on inferCNV clusters, transcriptomic clusters, and mitochondrial SNV clusters.

## Acknowledgements

The authors thank Eric Lam of The Garvan Weissman Centre for Cellular Genomics for assistance with Fluorescent-Activated-Cell-Sorting and acknowledge Winnie Luu and Alejandro Rios Villamil from The Garvan Weissman Centre for Cellular Genomics for single cell capture and 10x library construction, Ramaciotti Centre for Genomics (UNSW Sydney, Australia) for short read sequencing and SNP genotyping, Melissa Rapadas of The Garvan Institute of Medical Research for long read sequencing, Michael Li (UNSW) for assistance with data processing and analysis, and staff and donors of the Sydney Cord Blood Bank for providing cord bloods for research. Some biospecimens used in this research were obtained from the Health Precincts Biobank, UNSW Biospecimen Services, UNSW Sydney, Australia. The Biobank acknowledges the key contribution of NSW Health Pathology. This research was performed using the computational cluster Katana (https://doi.org/10.26190/669x-a286) supported by Research Technology Services at UNSW Sydney.

## Author contributions

JAIT, HRH, OS, AS, CJJ, JEP designed research; HRH, OS, XZ, SJ, MN, DH, DCW performed research; AS contributed vital analytical tools; JAIT, HRH, PLSB, OS, XZ, HMC, FCK, FY, DH, DCW, FV, MNP, YH, FZ, JEP analyzed and interpreted data; HRH, CJJ, JAIT, JEP wrote the manuscript.

## Competing Interests

The authors declare no competing interests.

## Notes

### Competing Interest Statement

The authors have declared no competing interest.

