## Supplemental Figures for "Single cell multiomics reveal clonal and functional dynamics of MDS stem/progenitor cells during hypomethylating therapy"

*Running title: Longitudinal changes in sub-clonal AZA response*

Julie A.I. Thoms[1,†,*], Henry R. Hampton[1,†], Priscilla L.S. Boon[2], Olivia Stonehouse[1], Xiaoheng Zou[1], Hoi Man Chung[3], Forrest C. Koch[4], Feng Yan[1,5], Swapna Joshi[1], Mary N.T. Nguyen[1], Dorothy Hung[6,7], Dale C. Wright[6,7], Fatemeh Vafaee[4,8], Mark N. Polizzotto[9], Alexander Swarbrick[10,11], Yuanhua Huang[3, 12], Christopher J. Jolly[1], Fabio Zanini[2,13,14,*], and John E. Pimanda[1,2,15, *].

[†] Equal contributions

**List of Supplemental Tables (see accompanying excel file)**

Supplemental Table 1: Patient characteristics

Supplemental Table 2: Single cell genotyping

Supplemental Table 3: SNP/CMA arrays

Supplemental Table 4: Sample pooling strategy

Supplemental Table 5: CITEseq antibodies

Supplemental Table 6: Primer sequences

Supplemental Table 7: Flow cytometry antibodies

Supplemental Table 8: Pool specific thresholds

Supplemental Table 9: Number of cell analyzed per patient and timepoint

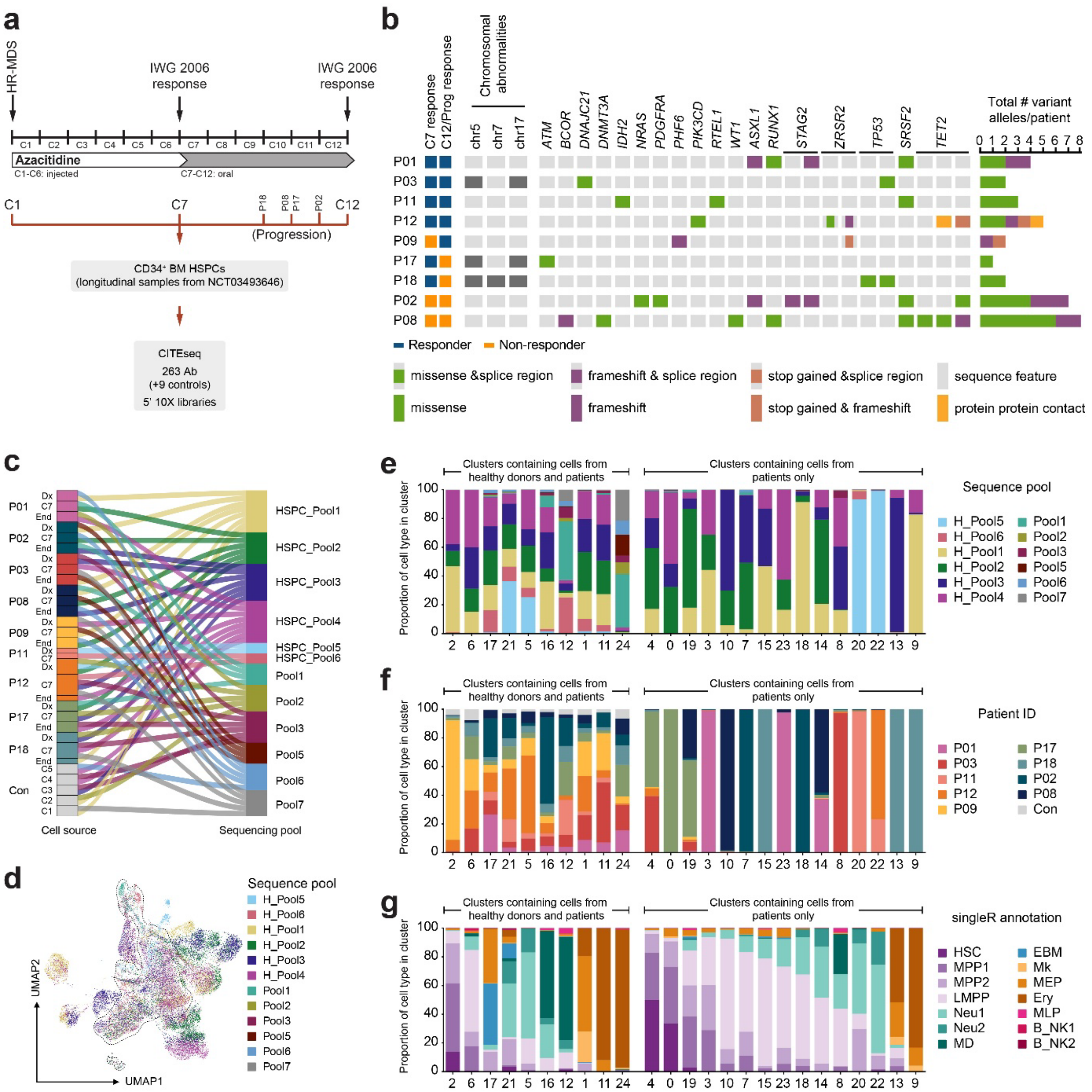


**Supplemental Figure 1**

**Supplemental Figure 1: MDS HSPCs contain transcriptomic clusters shared with healthy donors as well as patient-restricted transcriptomic clusters**

**a)** Schematic of clinical trial NCT03493646 showing clinical response and sample collection timepoints. **b)** Genetic features of HR-MDS patients at diagnosis. **c)** Sample pooling strategy. **d)** UMAP embedding coloured by capture pool. Dashed lines indicate 95% kernel density contours of UMAP coordinates for the shared clusters. **e)** Barplot showing proportion of cells in each leiden cluster derived from each capture pool. **f)** Barplot showing proportion of cells in each leiden cluster derived from each patient or from healthy donors. **g)** Barplot showing proportion of cells in each transcriptomic cluster assigned to each of 10 cell types by SingleR (Zhang et al Dev Cell 2022).

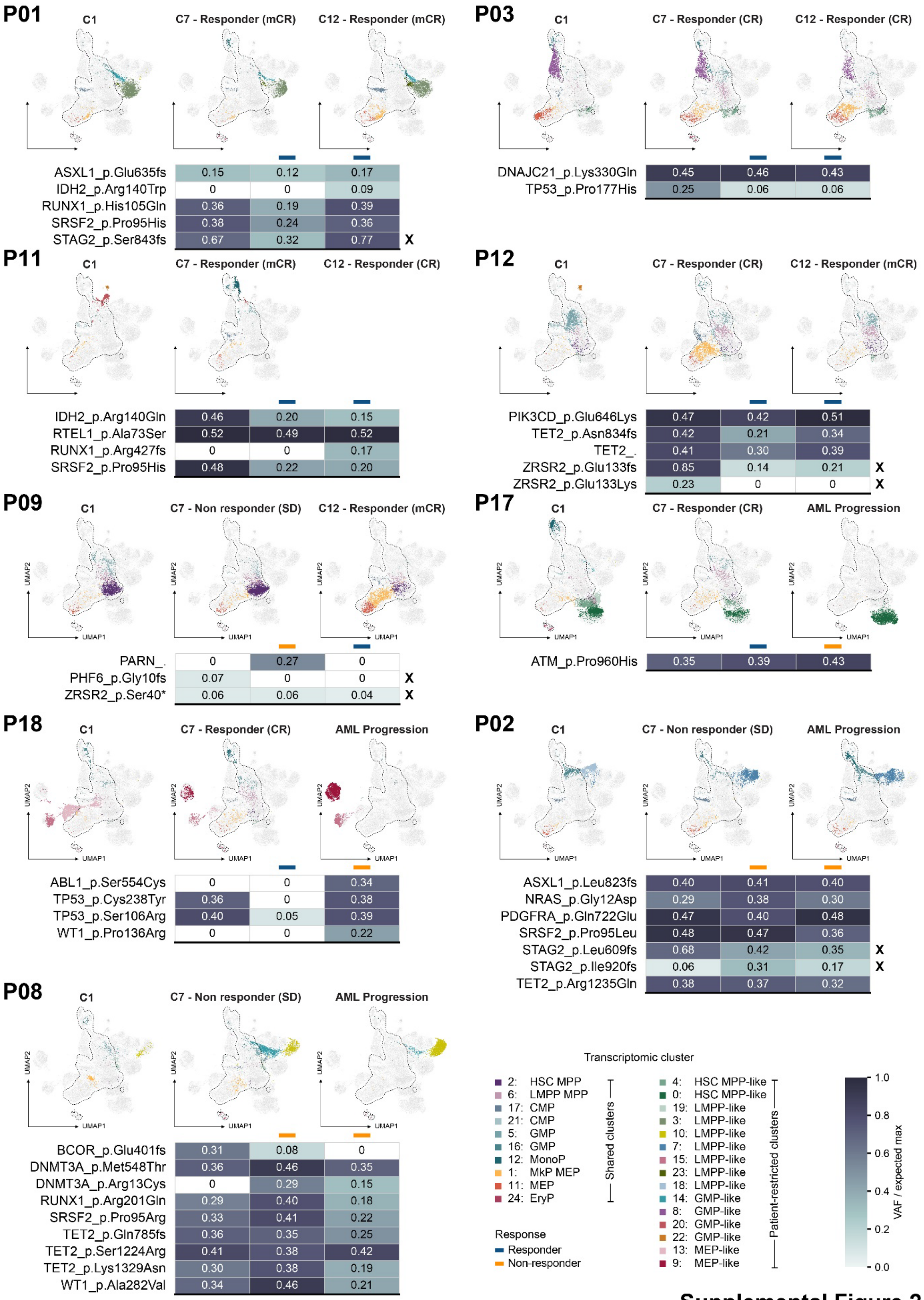


**Supplemental Figure 2**

**Supplemental Figure 2: Cluster and variant allele fraction changes in individual patients over the course of treatment**

UMAP embeddings highlighting shared and patient-restricted clusters at diagnosis (C1), following 6 treatment cycles (C7) and at end of treatment (C12 or Prog) for each patient. Dashed lines indicate 95% kernel density contours of UMAP coordinates for the shared clusters. Heatmaps show variant allele fraction (VAF) in BM MNCs from corresponding timepoints. Somatic variants with a detected VAF of at least 5% at any timepoint are shown. Number indicates measured VAF while heatmap shows detected VAF as a proportion of the expected maximum (expected maximum = 0.5 except for X-linked genes in male patients where expected maximum = 1; relevant variants are marked with an X). 0 indicates variant not detected at that time point. Dashed lines indicate 95% kernel density contours of UMAP coordinates for the shared clusters.

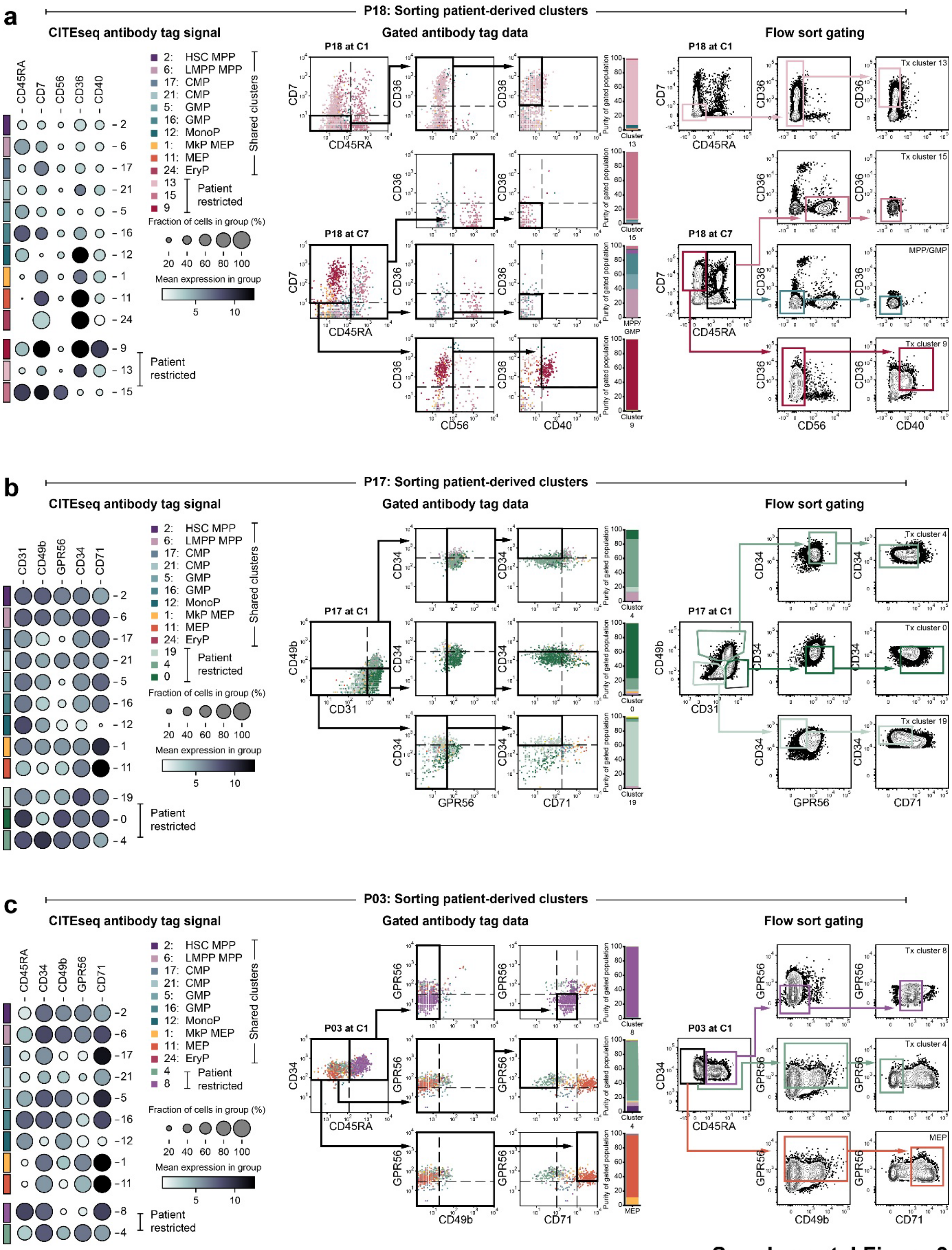


**Supplemental Figure 3**

**Supplemental Figure 3: Sorting cells from patient-restricted clusters**

**a)** P18, **b)** P17, **c)** P03. **a-c)** *Left:* Level and frequency of selected CITEseq antibody signal in shared and patient-restricted clusters. *Centre:* Gating scheme based on antibody tag data. *Right:* Flow sort gating used to prospectively isolate specific transcriptomic clusters.

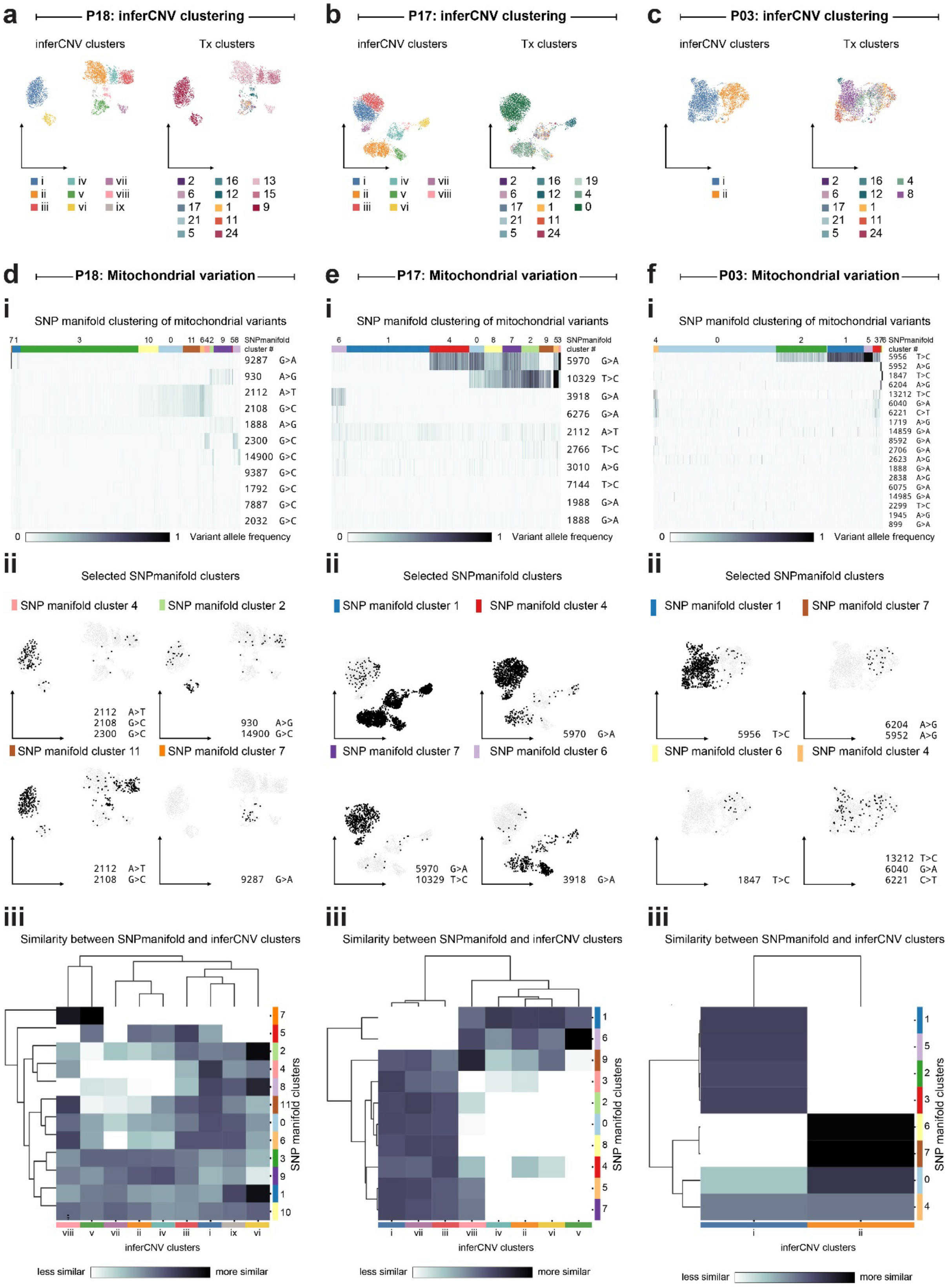


Supplemental Figure 4

**Supplemental Figure 4: Mitochondrial variants support inferred clonal relationships between transcriptomically defined patient-restricted clusters**

**a-c)** UMAP projections based on inferCNV profiles coloured by (left) inferCNV-based clusters and (right) transcriptomic clusters for **a)** P18, **b)** P17, **c)** P03. **d-f)** Analysis of mitochondrial variation **i)** SNP manifold clustering of mitochondrial variants based on mitochondrial variants detectable in 5' 10X data. **ii)** UMAP projections based on inferCNV profiles highlighting cells within selected discriminating SNP manifold mitochondrial clusters. **iii)** Similarity between SNP manifold mitochondrial clusters and inferCNV clusters in **d)** P18, **e)** P17, **f)** P03.

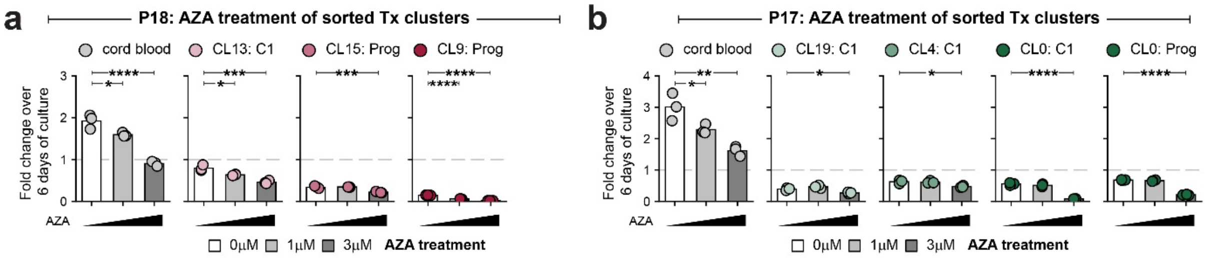


**Supplemental Figure 5**

**Supplemental Figure 5: Cells from sorted transcriptomic clusters have distinct behaviour in an in vitro co-culture system**

**a-b)** Fold expansion of total cells from **a)** P18 and control cord blood cells and **b)** P17 and control cord blood cells. Grey dashed line indicates initial cell seeding. Statistical comparisons are one-way ANOVA with Tukey's test for multiple comparisons, only comparisons to vehicle are shown. * indicates $P<0.05$, ** $P<0.01$, ***$P<0.001$, ****$P<0.0001$.
